# Deep learning algorithms reveal a new visual-semantic representation of familiar faces in human perception and memory

**DOI:** 10.1101/2022.10.16.512398

**Authors:** Adva Shoham, Idan Grosbard, Or Patashnik, Daniel Cohen-Or, Galit Yovel

## Abstract

Recent studies show significant similarities between the representations humans and deep neural networks (DNNs) generate for faces. However, two critical aspects of human face recognition are overlooked by these networks. First, human face recognition is mostly concerned with familiar faces, which are encoded by visual and semantic information, while current DNNs solely rely on visual information. Second, humans represent familiar faces in memory, but representational similarities with DNNs were only investigated for human perception. To address this gap, we combined visual (VGG-16), visual-semantic (CLIP), and natural language processing (NLP) DNNs to predict human representations of familiar faces in perception and memory. The visual-semantic network substantially improved predictions beyond the visual network, revealing a new visual-semantic representation in human perception and memory. The NLP network further improved predictions of human representations in memory. Thus, a complete account of human face recognition should go beyond vision and incorporate visual-semantic, and semantic representations.

## Introduction

In recent years, face recognition deep neural networks (DNNs) have closed the gap with humans, reaching human-level performance ^1,2^. Furthermore, recent studies reveal remarkable similarities between the representations of faces generated by these face-trained deep convolutional neural networks (DCNN) and humans’ neural and cognitive face representations (For review see: ^3,4^). For example, Grossman and colleagues (2019) revealed that the neural responses to faces in high-level visual cortex were well predicted by the representations of face-trained DCNN. Dobs and colleagues (2022) revealed brain-like functional specialization for faces in DCNNs optimized for the classification of objects and faces. Abudarham and colleagues (2021) showed that face-trained but not object-trained DCNNs are sensitive to the same view-invariant, critical facial features used by humans for face recognition. Recent studies have also shown that face-trained DCNNs generate a face inversion effect, similar to the well-established effect shown in humans ^8–10^. Finally, face recognition DCNNs show better classification for the race of faces they are trained with ^11–13^, similar to the other-race effect, which has been extensively investigated in humans ^14,15^. Thus, DNNs offer a promising method for exploring the nature of human face representations.

Despite these remarkable similarities between humans and face recognition DNNs, these studies primarily focus on perceptual matching of unfamiliar faces. This line of work overlooks the primary objective of human face recognition, which is the recognition of familiar faces. As social creatures, humans’ main goal is to identify the faces of individuals they socially interact with ^16,17^. Classification of unfamiliar faces, which DCNNs are optimized to excel in, hardly ever occurs in our daily life, except in the realm of suspect recognition or security checks. Whereas two recent studies did propose a DCNN model of human familiar face recognition, they only concerned the visual representation of familiar faces in perception ^18,19^. However, cognitive and neural models of face recognition indicate that familiar faces differ from unfamiliar faces in at least two fundamental ways that are not considered by current face recognition DCNNs. First, familiar faces are not pure visual representations but are inherently associated with semantic information ^20–22^. Second, successful recognition of familiar faces depends on matching a perceptual representation to a representation in memory ^21^. Nonetheless, the nature of the representation of familiar faces in memory, which is critical for recognition, were not considered so far by face recognition DCNNs.

To close this gap between human and machine face recognition, in the current study we employed two recent DNNs to represent semantic aspects of familiar faces. CLIP (Contrastive Language Image pre-training^23^) is a recent DNN that was trained to pair images and text based on internet webpages. Unlike standard visual DNNs that link images to arbitrary labels, the text caption of an image in a webpage is semantically related to the image. Whereas CLIP is designed for general object recognition, it is also trained on celebrity faces and their text caption. Thus, CLIP offers an opportunity to explore a novel visual-semantic representation of familiar faces, which has not been explored yet. In addition, we examined whether this visual-semantic representation is distinct from pure visual and pure semantic representations. To that effect, we used a recent natural language processing (NLP) algorithm (SGPT ^24^), to extract pure semantic representation of familiar identities based on their textual description from Wikipedia. This enabled us to examine whether visual-semantic and semantic DNNs account for additional information in the representation that is generated by humans for familiar faces, beyond the visual information that has been so far investigated with standard visual DCNNs trained to classify face images (e.g., VGG-16 ^25^).

In the current study, we used visual (VGG-16), visual-semantic (CLIP) and semantic (SGPT) DNNs to predict human representations of familiar faces in perception and memory. To study human face representations, human participants were asked to rate the visual similarity of famous faces identities based on their face images or the reconstruction of their visual appearance from memory, based on their names (Figure 2A). These similarity measures were used to construct the representational geometry of familiar faces in perception and memory (Figure 2B-C). We used the representational geometries of the same identities based on visual (VGG-16), visual-semantic (CLIP) and semantic (SGPT) DNNs (see Figure 1) to predict human representations in perception and memory (Figure 2D-E). In particular, we assessed whether visual-semantic and semantic DNNs improve predictions of human representations beyond the commonly used visual face recognition DCNN, and what is the relative contributions of these different representations in perception and memory. This will enable us to assess whether visual-semantic and semantic information contributes to the representation of faces in perception and memory beyond pure visual information.

**Figure 1:**
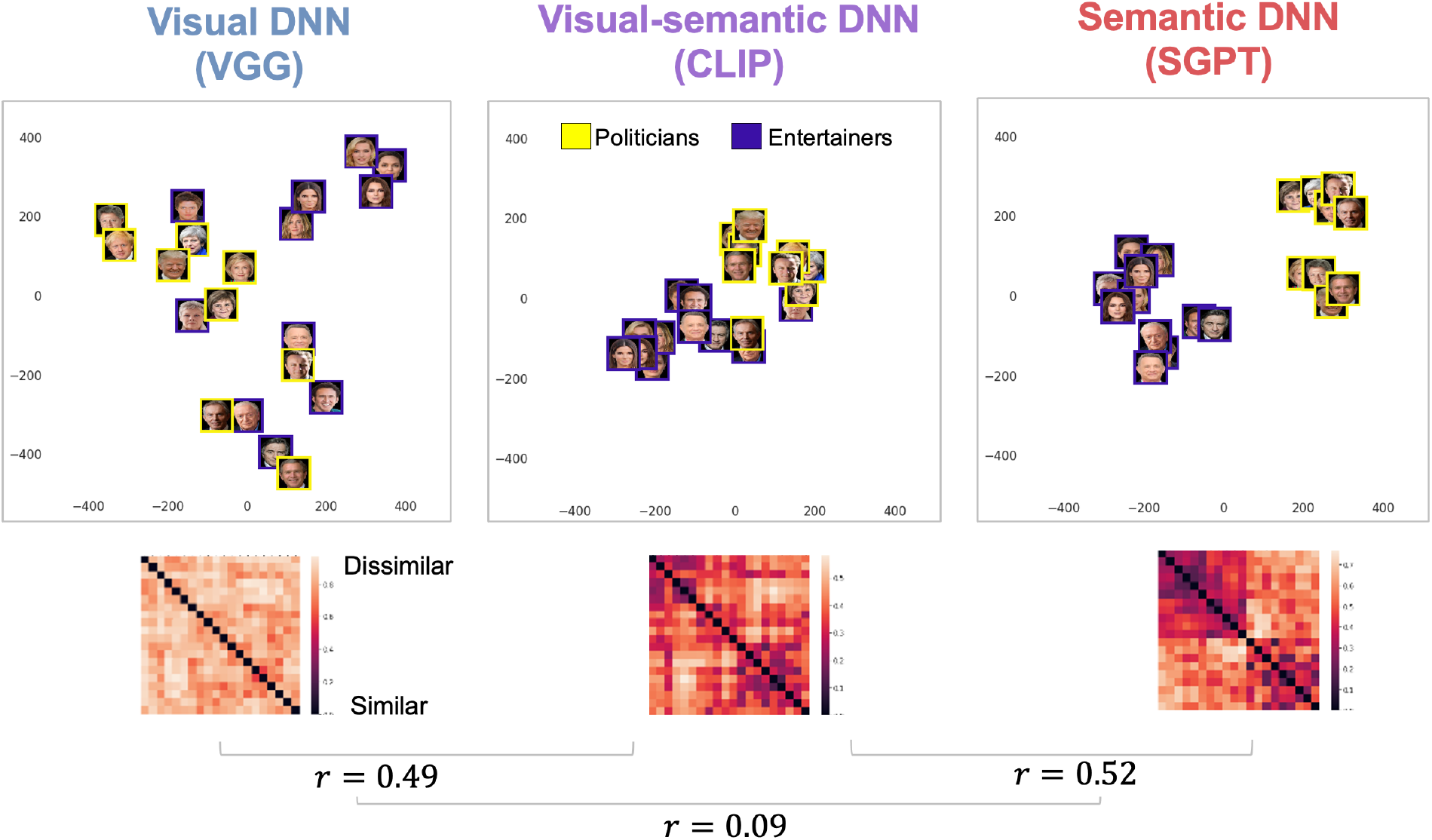
A 2-D visualization of the representational geometry of familiar faces based on the RDMs of their embeddings in visual (VGG), visual-semantic (CLIP) and semantic (SGPT) DNNs. VGG and CLIP representations are based on images and SGPT representation is based on text from Wikipedia of the same familiar identities. The SGPT t-SNE displays face images but is only based on the text that describes these identities. The CLIP visual-semantic representation was correlated with both the pure visual (VGG) and pure semantic (SGPT) representations, whereas no correlation was found between the visual (VGG) and semantic (SGPT) representations.

**Figure 2:**
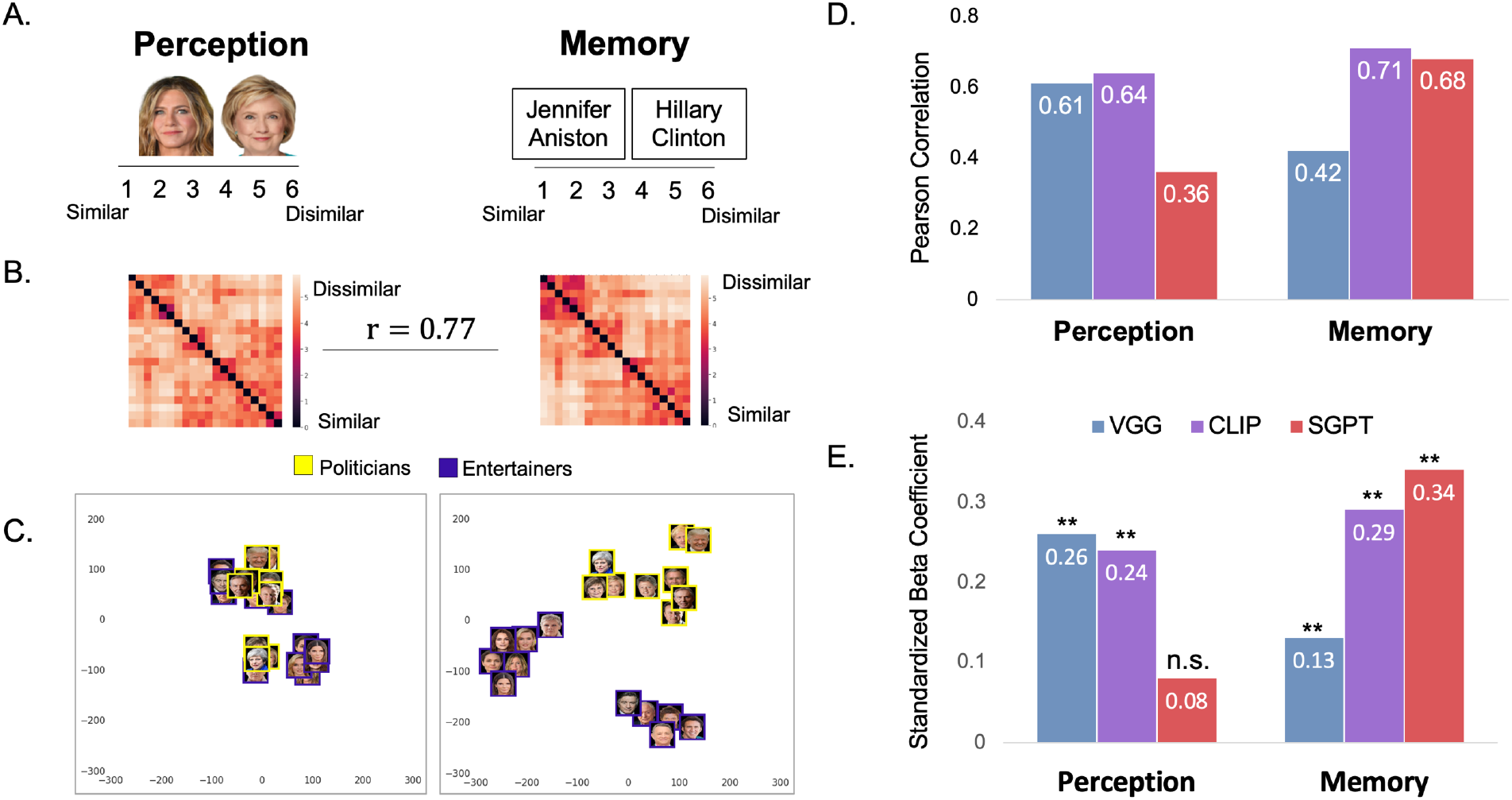
A. Participants rated the visual similarity of face images (perception) or the reconstruction of the appearance of these faces from memory based on their names (memory). B. RDMs based on human visual ratings in perception and memory. The representations in perception and memory were highly correlated. C. A t-SNE visualization of the representational geometry of the familiar faces, showing semantic influence on the visual representation in memory but less in perception. D. Correlations between the RDMs based on embeddings of the same identities in visual (VGG-16), visual-semantic (CLIP) and semantic (SGPT) DNNs with the visual representations in perception and memory. E. Results of the multiple linear regression model with the three DNNs as predictors of human visual perception and visual memory show that visual (VGG-16) and visual-semantic (CLIP) DNN each uniquely predict human representations in perception, with no contribution from a pure semantic (SGPT) DNN; The representation of familiar faces in human memory is predicted by visual-semantic (CLIP) and semantic (SGPT) DNNs, and to a lesser extent the visual (VGG) DNN (see also Figure 3, right column). ** p < .001

## Results

### The representational geometry of familiar faces by visual, visual-semantic, and semantic DNNs

We examined the representational geometry of 20 internationally famous identities, 10 politicians and 10 entertainers. To obtain visual-semantic representations of familiar faces, we selected only identities that were included in the training set of CLIP (see methods section and supplementary Figure 1). We measured the cosine distance between the embeddings of the face images in VGG-16 and CLIP as well as the SGPT embeddings of the first paragraph of their Wikipedia text (see Supplementary Table 1 for the Wikipedia text). Figure 1 shows the representational dissimilarity matrices (RDMs) and a t-SNE 2D visualization ^26^ of the geometry of the identities according to each of the three DNNs. The identities were clustered by their occupation in the semantic DNN (SGPT), and the visual-semantic DNN (CLIP), but not the visual DNN (VGG). The correlations between the RDMs of the different algorithms show that the visual-semantic representation generated by CLIP was correlated with both the pure visual representation of VGG (r = 0.49) and the pure semantic representation of SGPT (r=0.52), whereas the visual (VGG) and semantic (SGPT) representations were not correlated (r=0.09).

### Visual-semantic and semantic DNNs significantly predict human representations of familiar faces beyond the standard visual face recognition DNN

To assess the nature of the representations of familiar faces in perception and memory, human participants rated the visual similarity of the same 20 familiar identities based on their images (perception) or the reconstructions of their facial appearance from memory based on their names (memory) (Figure 2A) (see Supplementary Figure 2 for reliabilities of human similarity ratings). We first computed the correlation between the RDMs of visual similarity ratings in perception and memory (Figure 2B). The correlation between visual similarity ratings in perception and memory was very high (r = 0.77), indicating that participants generated a visual image of the familiar faces in memory. We used t-SNE to visualize the representational geometry of the faces in perception and memory based on their RDMs. As can be seen in Figure 2C, the identities are clustered based on their occupation (politicians or entertainers) in memory, but this arrangement is less apparent in perception. Thus, although the visual representations in perception and memory are highly correlated, they appear to differ in the relative contribution of visual and semantic information.

To quantify the contribution of visual and semantic information to humans’ representations of familiar faces, we examined whether the representations of familiar faces in human perception and memory are correlated with the representations of the same identities in visual (VGG-16), visual-semantic (CLIP), and semantic (SGPT) DNNs. The correlations reported here are based on the representations in the penultimate/output layer, which is used for classification of each network (see Supplementary Figure 3 for the correlations of human perception and memory with VGG and CLIP across all their layers). We computed the correlations between the RDMs of human similarity ratings of images (perception) and names (memory) with the RDMs of the same identities of VGG-16, CLIP and SGPT (see Supplementary Figure 4 for the full correlation matrix). As shown in Figure 2D, human visual perception is correlated with a pure visual (VGG-16) (r = 0.61) and a visual-semantic (CLIP) (r = 0.64) DNNs and less so with a pure semantic DNN (SGPT) (r = 0.36), whereas human visual memory is more correlated with visual-semantic (CLIP) (r = 0.71) and semantic (SGPT) (r = 0.68) DNNs than a pure visual (VGG-16) DNN (r = 0.42). Thus, visual-semantic and semantic DNNs account for significant proportions of variance in human representations of familiar faces.

Given that CLIP is correlated with both the pure visual (VGG) and pure semantic (SGPT) representations, it is possible that it does not contribute any unique information beyond the pure visual and semantic codes. To address this question, we next examined the unique contribution of each of the DNNs to human representations in perception and memory with a linear regression model. We started with VGG as a sole predictor of human visual representations, as this is the network that has been typically used to predict human perception in previous studies of face recognition ^5,18,27^. Then we examined whether CLIP and then SGPT account for an additional proportion of variance in human perception and memory beyond the pure visual face recognition DNN (VGG-16). The results of these analyses are reported in Figures 2E and Figure 3.

**Figure 3:**
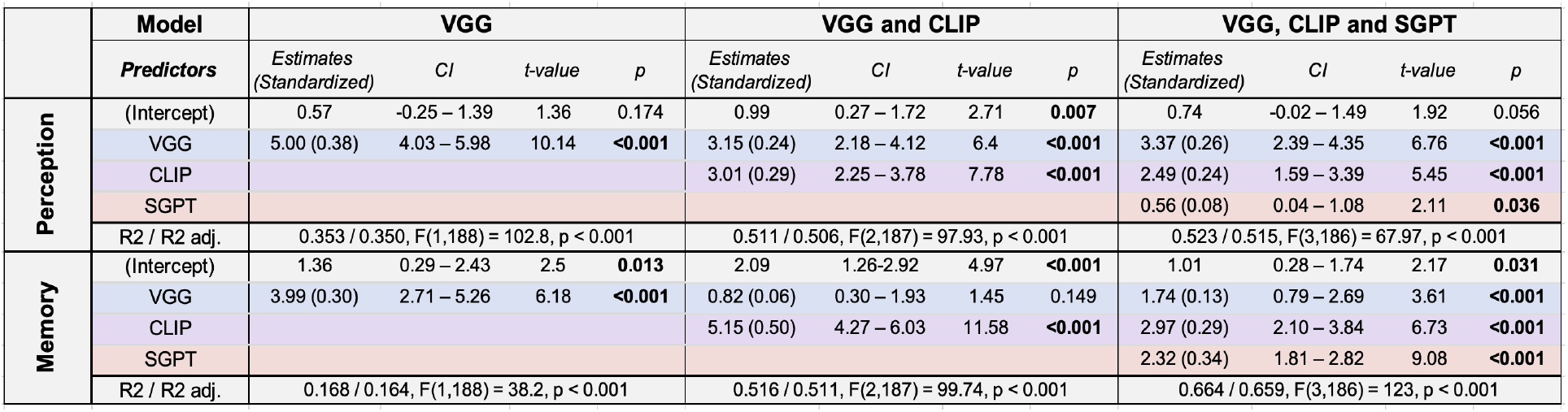
A summary of results of linear regression models in which VGG (left), VGG and CLIP (middle) and VGG, CLIP and SGPT (right) are used as predictors of the representations of faces in human perception (top) and human memory (bottom). The three deep learning algorithms accounted for 53% of the total variance in human perception and 67% of the total variance in human memory.

A linear multiple regression analysis showed that VGG alone accounted for 35% of the total variance in human visual perception and 17% in visual memory. Adding CLIP significantly improved predictions, with both DNNs accounting for 51% of the total variance in visual perception and 51% of the total variance of visual memory. Finally, adding SGPT as a third predictor significantly improved the prediction of the representations in human visual memory, accounting for 67% of its total variance, but did not account for additional variance in human visual perception beyond VGG and CLIP.

To assess whether the semantic representations generated by SGPT are valid models of human semantic representations, we also asked human participants to rate the semantic similarity of the same identities (see methods). Participants were asked to ignore the visual appearance of the identities and judge them only based on their biographical information. Figure 4 shows the correlations between the three DNNs and human semantic ratings (see Supplementary notes and Supplementary Figures 5 and 6 for additional information about the semantic representations). Notably, the correlation between SGPT and human semantic ratings was very high (r = 0.84) indicating that SGPT representations almost fully account for human semantic representations. Figure 4 also shows the correlations of the three DNNs with visual perception (green) and memory (gray) (also displayed in Figure 2D in blue (perception) and red (semantic)), so they can be directly compared to the correlations with human semantic ratings (yellow).

**Figure 4:**
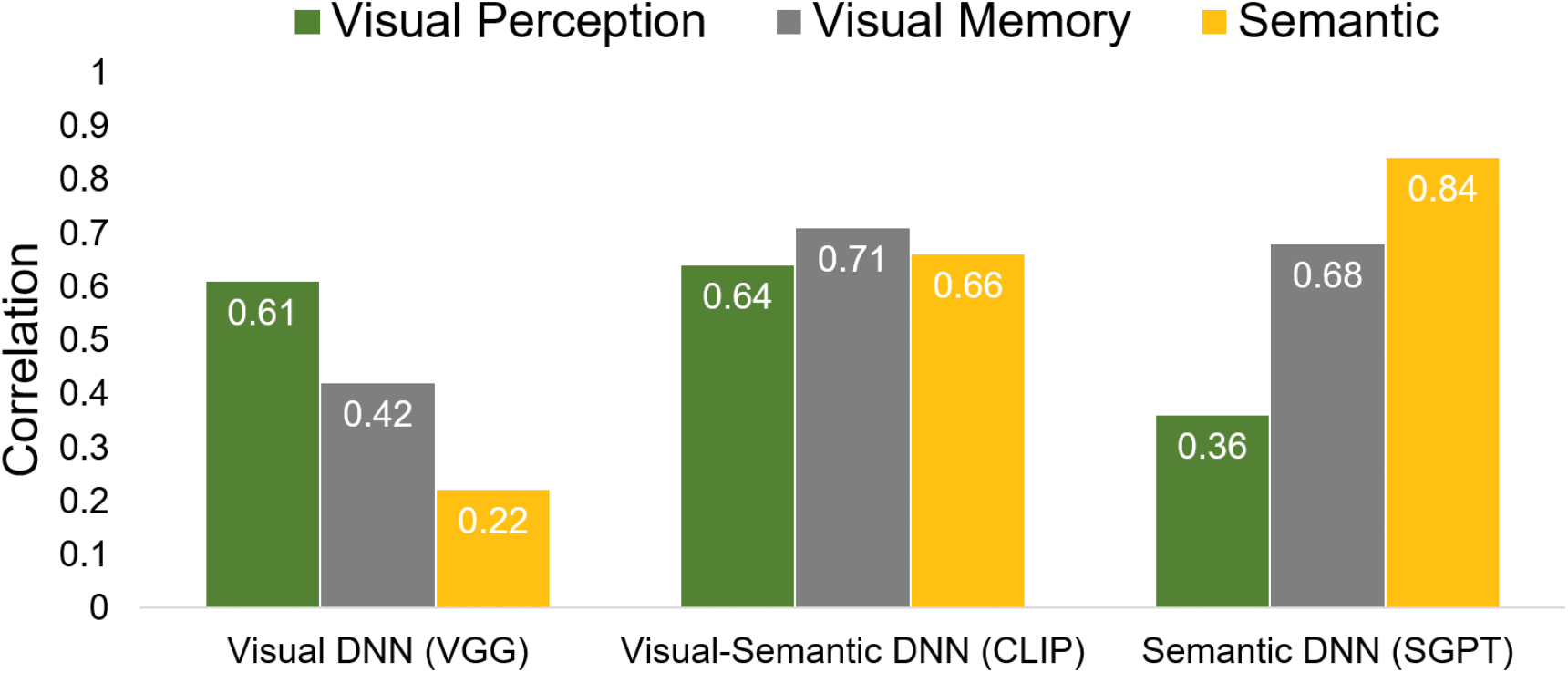
The correlations between visual (VGG), visual-semantic (CLIP) and semantic (SGPT) DNNs and human semantic similarity ratings (yellow), show high correlations between human and DNN semantic representations. For comparison, the figure also shows correlations with human visual perception and visual memory (same data presented in Figure 2D).

Taken together, our findings show that visual and visual-semantic DNNs account for over 50% of the variance in human visual perception and visual memory. A pure semantic DNN increases the total explained variance in human memory to 67%. Overall, these findings show that visual-semantic and semantic DNNs significantly improve the prediction of human representations of familiar faces, beyond the pure visual algorithm that has been so far used to model human face representations ^3,4,6–8,10,11,28–30^.

CLIP was a strong predictor of both visual perception and visual memory. Current cognitive and neural models of face recognition presume that familiar faces are represented by distinct visual and semantic codes ^20,21,31^. The visual-semantic representation that is generated by CLIP accounts for an additional variance that is not explained by pure visual and semantic models. This suggests a new type of visual-semantic code that has not been considered in models of human face recognition so far. To further examine the nature of the representation that is generated by a visual-semantic algorithm and how it may differ from the representation that is generated by a pure visual representation, we used StyleGAN, a generative adversarial network ^32,33^, to generate faces based on their embeddings in VGG and CLIP (see Methods). Figure 5A shows an example of the faces of Hillary Clinton and Jennifer Anniston that were generated by StyleGAN based on the embeddings of their identities in VGG and CLIP (see methods and supplementary notes for the procedure used to generate the faces; See supplementary Figure 7 for the VGG and CLIP-generated faces of all 20 identities). These AI-generated images provide us with a new way to assess the similarity between human perception and memory and the representations generated by VGG and CLIP. To that effect, we asked a new group of human participants to rate the visual similarity of the VGG- or CLIP-generated faces. We then used these RDMs to predict the representations of the original identities in human perception and memory. Figure 5A shows the RDMs of human similarity ratings of the VGG- and CLIP-generated faces. Figure 5B shows that RDMs based on human similarity ratings of VGG- and CLIP-generate faces predicted human visual perception of images of the original identities (R^2^ = 0.57, p < 0.001), whereas human visual memory was predicted by human ratings of CLIP-generated faces but not VGG-generated faces (R^2^ = 0.49, p < 0.001).

**Figure 5:**
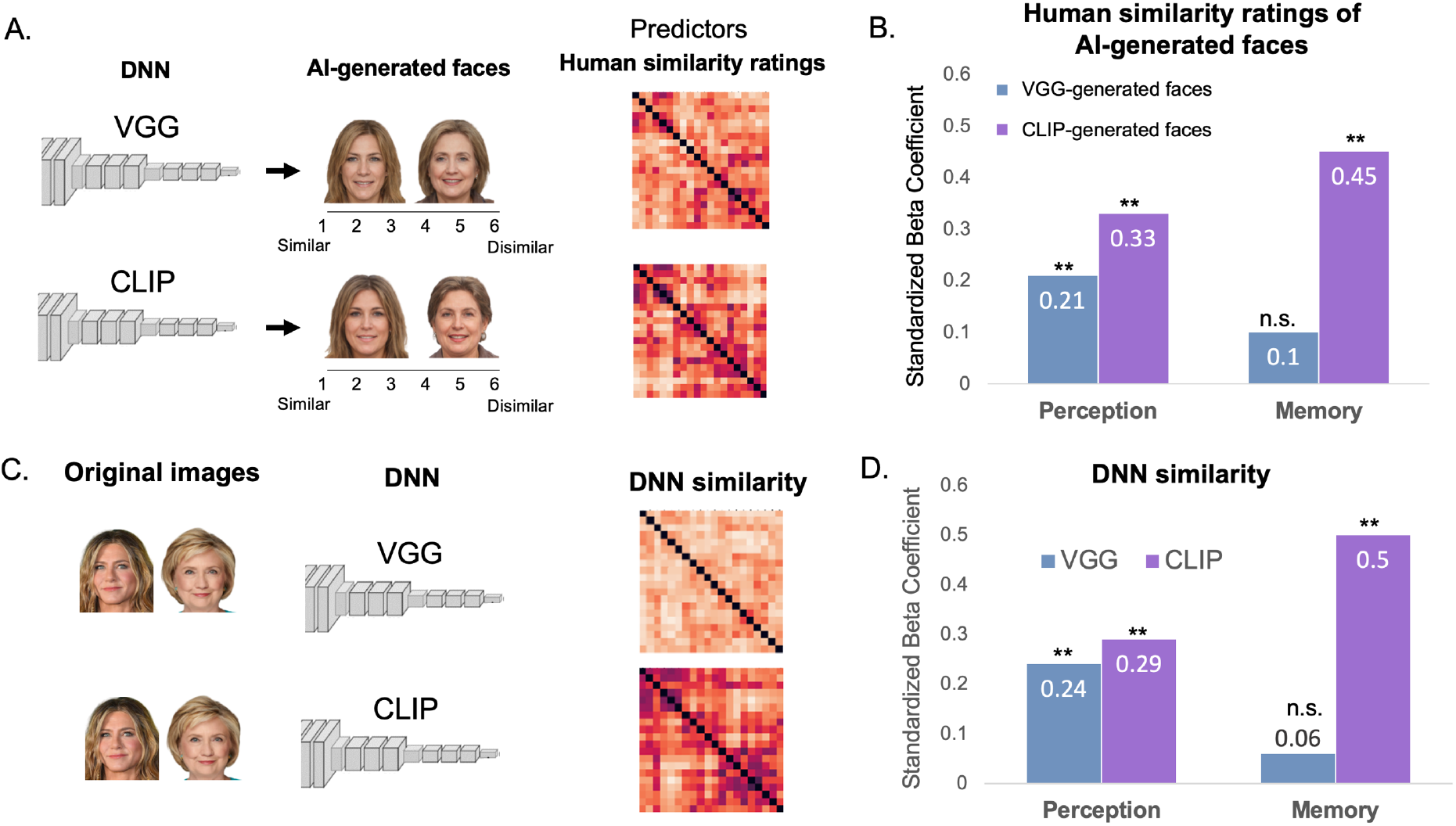
A. RDMs of human similarity ratings of VGG-generated faces (top) and CLIP-generated faces (bottom). B. Human similarity ratings of VGG-generated and CLIP-generated faces were used in a linear regression to predict the similarity rating of the original faces of the same identities in perception (images) or memory (names). C. The RDM of VGG (top) and CLIP (bottom) according to the embeddings of the original images in VGG and CLIP (see also Figure 2B) D. The RDMs based on VGG and CLIP embeddings were used as predictors of the representations in perception and memory (Figure 3 middle column) show similar findings indicating a strong correspondence between human and algorithms representations. ** p < .001

The same regression that is based on the RDMs of VGG and CLIP embeddings of the original images (see Figure 3, middle column, Figure 5D) revealed very similar results. These results indicate that human ratings of VGG- and CLIP-generated faces, closely correspond with the networks’ embeddings of the original identities (see Supplementary figure 8 for the correlations between them). Overall, these findings indicate that the visual representations of familiar faces are shaped by their semantic content. These representations can be modelled by an algorithm that learns to classify images by linking them with their non-visual semantic information.

## Discussion

Face recognition DCNNs show remarkable similarities with human cognitive and neural representations ^3,4,6–8,10,11^. These algorithms are trained on face images and therefore exclusively rely on visual information. Here we proposed that human face recognition goes beyond vision as it primarily involves the recognition of familiar faces, which are represented by visual and semantic codes ^20,21,31^. Furthermore, human familiar face recognition involves matching a representation of an input image in perception with a visual representation in memory. Nonetheless, recent studies that have shown correspondence between human and face recognition DCNNs were only concerned with the perceptual representation that are generated for the input images ^5,6,27^. Thus, in the current study we examined whether recent visual-semantic and semantic DNNs may account for additional variance in the representations of familiar faces in perception and memory, beyond pure visual information. Our study reveals the following novel findings: First, a visual-semantic (CLIP) algorithm uniquely accounted for the visual representation of faces in human perception, beyond the standard visual face recognition algorithm (VGG-16). Second, the representation in memory, which has not been explored before, is mainly predicted by visual-semantic and semantic DNNs. We therefore conclude that visual information alone does not account for human representations of familiar faces, but there is an additional significant contribution of semantic information. Moreover, we revealed a new visual-semantic representation in human perception and memory that was unknown so far.

Our findings do not only advance our understanding of the type of DNNs that best predict human representations, but also provide new insights on current cognitive models of human face recognition. A central assumption of current face recognition models is that the visual and semantic codes of familiar faces are mediated by distinct cognitive and neural mechanisms ^20,21,31^. According to a seminal model of human face recognition ^21,31^, familiar faces are represented in a face recognition unit (FRU) that is purely visual. This visual representation is linked to semantic information in a person identity node (PIN). A neural model of familiar face recognition by Gobbini and Haxby (2007) further suggests that visual information from faces is represented in the core face-selective areas in high-level visual cortex, whereas semantic information is represented in the extended non-visual brain regions, including the limbic system, precuneus and the anterior temporal lobe.. The visual-semantic representation that is generated by CLIP suggests a new module where visual and semantic information may be integrated, which is separated from pure visual and pure semantic representations. This visual-semantic representation that we found both in human perception and human memory may reflect an intermediate representation that links between the pure visual information that is encoded by the visual system and the pure semantic information about the image that is represented in memory. A possible neuroanatomical location of this type of visual-semantic representation is the perirhinal cortex which integrates visual and semantic information ^34^. A recent study in macaques found distinct neural systems for personally familiar faces in the perirhinal cortex and temporal pole ^35,36^. These anterior temporal representations may be better predicted by CLIP than by current visual DCNNs that are trained on faces linked to meaningless, arbitrary labels.

The similarity between human mental representations and DNN’s representations is typically based on the distance between the embeddings of the images. Current face generator algorithms (StyleGAN) enable us to visualize the nature of these embeddings. This procedure provides us with a new way to assess this correspondence between human’s and DNNs’ representations, by asking human participants to rate the similarity between AI-generated faces and use these RDMs as predictors of human representations of the original images in perception and memory. The results of these analyses were remarkably similar to the predictions based on VGG and CLIP embeddings of the original images (Figure 5B, D). The representational geometry of faces was clustered by their occupation based on human ratings of CLIP-generated faces but not VGG-generated faces (Supplementary Figure 8). These findings further demonstrate the close correspondence between human and DNNs representational geometries, indicating that they can be used as valid models of human face representations.

Our findings show that even pure semantic information, extracted by NLP algorithms, can account for the representation of familiar faces in memory, beyond visual and visual-semantic information. These non-visual, semantic representations were based on a textual description of non-visual, semantic information about the familiar identities in Wikipedia and were indeed uncorrelated with their visual representations generated by VGG (Figure 1). Thus, even though participants were instructed to judge the visual similarity of the faces of the familiar identities, pure semantic information intruded the representation in memory, but not in perception. These findings show how DNNs enable us to isolate the contributions of visual, visual-semantic and semantic information, and this way uncover the content of the representations in perception and memory, which are hard to access and explore.

The main goal of machine learning algorithms is to successfully classify untrained examples. Accordingly, the performance of face recognition algorithms is measured by their classification of untrained faces. This operation is analogue to identity matching tasks with unfamiliar faces in humans ^2,7,19,37^. Whereas face-trained DCNN may reach and even surpass human-level performance on such identity matching tasks, human face recognition in daily life is rarely concerned with the discrimination of unfamiliar faces, which is the main task DCNNs are optimized to perform. Thus, a valid computational model of human face recognition should concern the type of operations that humans typically perform, which is the identification of socially relevant, familiar faces. Familiar faces are not pure visual representations but are inherently linked to unique semantic information. The representation that is generated by CLIP, which is trained to link images to their meaningful semantic text in a self-supervised manner may be a more suitable model that should be further explored in future studies that examine the neural and cognitive mechanisms of human face and object recognition.

In conclusion, human intelligence relies on cognitive operations that integrate different codes into a unified representation. Our findings show that even face recognition, which is typically considered a visual problem, is best predicted by visual and semantic DNNs. Deep learning algorithms enable us to examine the nature of these visual, visual-semantic, and semantic representations and their unique contributions to perception and memory in a way that was not possible before. The approach we used here is not limited to faces and may provide a new framework and the computational tools to model human representations of other categories and domains including familiar objects, places as well as sounds and voices. Our findings also inform machine intelligence in that they highlight the importance of multi-model systems that integrate sensory and semantic information. Future AI models of human cognition may also integrate emotional, and motor information as well as attentional and motivational factors that select behaviorally relevant information. This type of multi-system operation may underlie the efficient learning and adaptive behavior that is required to close the gap between computer and human general intelligence. Overall, our findings show how human and machine intelligence may inform and advance each other by offering new insights and approaches that are less evident when each system is studied alone.

## Methods

### Stimuli

Original faces: We selected 20 identities of international celebrities, 10 politicians and 10 entertainers, that were included in the CLIP training set (see supplementary Figure 1 for a complete list of the identities) and generate a visual-semantic representations of their face images. To select identities that CLIP was trained with, we examined whether face images of the selected identities could be correctly classified by CLIP without finetuning, following the zero-shot classification method proposed by Radford et al (2021). We measured the cosine similarity between the embeddings of names and images of these identities based on CLIP and selected only identities that the similarity between their image and their name was the highest relative to any of the other names (see supplementary Figure 1).

We then selected face images of these identities from a Google images search. The face images were colored images that included only the face of each identity. The background of the images was removed and all images were aligned to the same size. The names of the identities were presented on white background (see Figures 2A).

DNN generated faces: To generate faces, we used the SOTA generator for human faces – StyleGAN ^32,33^. Our goal was to transform VGG and CLIP embeddings to StyleGAN’s embeddings. To do so, we first trained a model to generate StyleGAN’s embeddings based on the DNN’s embeddings; the full training procedure is described in the supplementary notes. We used this model to generate images of the identities used in the experiment based on VGG and CLIP representations (Figure 5A). To reduce the effect of noise found in the DNNs’ embeddings of a single image, which can be transferred to the generated images, we used the DNNs’ representations of 20 different images of each identity and calculated their mean StyleGAN’s embedding. Using this mean we generated each image.

#### Deep neural networks

##### Face-trained VGG-16

To obtain a pure visual representation of the face images we used the VGG-16 algorithm ^25^ and trained it to classify 8749 images of faces from the VGGFace2 dataset ^39^. All images were first aligned using the MTCNN algorithm ^40^. Training and image preprocessing followed the procedure from Parkhi et al (2015) with the following changes: Images were normalized according to *μ* = [0.5,0.5,0.5],σ = [0.5,0.5,0.5], the training was done on batches of 128 images, for 120 epochs of 1000 batches and after 100 epochs, learning rate was reduced to 1e-3. The network’s performance was measured on a face verification task consisting of 6000 pairs of face images from the Labeled Faces in the Wild benchmark ^42^. The network’s verification performance was 97%. To extract the representation of each face image of the study stimuli, the images were first aligned using the MTCNN algorithm ^40^. We then extracted the embeddings based on the feature vector representation in the penultimate (fc7) layer of the network and computed the similarity between each face pair based on the cosine distance between this feature vectors. In Supplementary Figure 3 we also show these correlations with all other layers of the network.

##### CLIP (Contrastive Language-Image pre-training)

CLIP is trained to create similar representations for images and their text caption based on 400M images from the internet ^38^. We extracted the embeddings of each face image based on the output layer of trained ViT-B/32 architecture. In supplementary Figure 3 we also show these correlations with all other layers of the network. We computed the similarity between each face pair based on the cosine distance between these representations. All face images were aligned using the MTCNN algorithm ^40^, and then preprocessed according to the values supplied by OpenAI’s implementation ^23^.

##### SGPT

GPT Sentence Embeddings for Semantic Search is a recent natural language processing (NLP) algorithm that is first pre-trained to predict the next word in a sentence similar to other NLP algorithms and use contrastive fine-tuning to create similar representations for pairs of sentences that describe the same content ^24^. We extracted the embeddings of the text of the first paragraph in Wikipedia of each identity based on the 1.3B parameters bi-encoder’s output layer and compute the similarity between each identity pair based on the cosine distance between these representations. Supplementary Table 1 shows the first paragraph in Wikipedia that was used for each identity.

#### Human similarity ratings

##### Participants

A total of 80 participants were recruited for this study from the Prolific platform. 20 participants were assigned to each of the four experimental conditions: Visual similarity based on images or names and semantic similarity based on images or names. Two participants were excluded from the analysis (1 participant from the visual memory condition, and 1 participant from the semantic (images) condition) because they were not familiar with 30% or more of the presented identities, which resulted in a total of 78 participants (mean age 30 (sd = 2.9), 60 females). The participants were paid 4 GBP for their participation in the experiment (8 GBP/hour). They gave informed consent prior the experiment. The study was approved by the ethics committee of Tel Aviv University.

A total of 40 additional participants were recruited (mean age 31 (sd = 2.7), 30 females) from the prolific platform. 20 participants were assigned to each of the two experimental conditions: visual similarity ratings according to VGG generated images or CLIP generated images. The participants were paid 4 GBP for their participation in the experiment (8 GBP/hour). They gave informed consent prior the experiment. The study was approved by the ethics committee of Tel Aviv University.

##### Procedure

Participants rated the visual or semantic similarity of all possible pairs of the 20 identities.

###### Visual similarity rating

Each trial presented one pair of images or names of two different identities, and the participants were asked to rate the visual similarity of their faces either based on the images (perception condition) or the reconstruction of their faces from memory, based on their name (memory condition). In the memory condition, we emphasized that similarity should be based on the visual appearance of the face based on their memory. The image/name pairs were presented on the screen one at a time, above a similarity scale (1 (very similar) - 6 (very dissimilar)) until response. The participants selected the similarity score with the mouse. The next pair was presented 1 second after their response. The participants had a forced break for a minimum of 10 seconds after 80 pairs were presented, and again after 160 pairs were presented. After they rated all 190 pairs, the participants were asked to indicate for each face/name whether they were familiar with it before the experiment. The experiment lasted about 30 minutes.

The same procedure was used to collect human similarity ratings of the DNN-generated faces.

###### Semantic similarity rating

Participants were asked to rate the similarity of the identities based on biographical or any other semantic information that they know about them. We emphasized that the similarity should not be based on visual appearance but only on semantic information.

#### Data analysis

##### Representational similarity matrices (RDMs) and regression analysis

We generated RDMs based on human similarity ratings averaged across participants and the embedding of the same identities in VGG-16, CLIP and SGPT. We also generated RDMs based on human similarity ratings averaged across all participants of the VGG and CLIP-generated images.

###### Human RDMs

We computed the averaged similarity ratings of each pair of faces or names across participants. We also computed the reliability between the participants’ ratings of the original stimuli in each condition (See supplementary Figure 2).

###### Deep neural network (DNN) RDMs

We measured the cosine similarity between the embedding of the same identities in the penultimate layer of VGG and in the output layer of CLIP, based on their images and in the output layer of SGPT based on the first paragraph of their Wikipedia textual description.

###### Correlation and regression analysis

We computed the correlations between human and DNN similarity ratings. To assess the unique contribution of the different algorithms to the variance in human similarity ratings, we used multiple linear regression in which VGG was the first predictor and then added CLIP and SGPT to assess the additional proportion of variance that each algorithm explained in the human data and its statistical significane.

To assess the similarity of human visual representations of the original stimuli and the DNN-generated images, we used multiple linear regression in which human visual similarity ratings of VGG- and CLIP-generated images were the predictors of human visual similarity ratings of the original face stimuli.

###### Representational geometry visualization

We used t-SNE, a nonlinear dimensionality reduction technique ^43^ for visualization purposes only. The correlational and regression analyses were based on the cosine similarity measures (RDMs).

## Acknowledgement

We thank Rafi Malach, Michael Gilead, Idan Blank and Daphna Shohamy for discussion of findings reported in this manuscript. This work was funded by an Israeli Science Foundation grant (ISF 917/2021) to GY.

## References

1. Taigman, Y., Yang, M., Ranzato, M. & Wolf, L. Deepface: Closing the gap to human-level performance in face verification. in Proceedings of the IEEE conference on computer vision and pattern recognition 1701–1708 (2014).

2. Phillips, P. J. et al. Face recognition accuracy of forensic examiners, superrecognizers, and face recognition algorithms. Proc. Natl. Acad. Sci. 115, 6171–6176 (2018).

3. O’Toole, A. J., Castillo, C. D., Parde, C. J., Hill, M. Q. & Chellappa, R. Face space representations in deep convolutional neural networks. Trends Cogn. Sci. 22, 794–809 (2018).

4. O’Toole, A. J. & Castillo, C. D. Face recognition by humans and machines: Three fundamental advances from deep learning. Annu. Rev. Vis. Sci. 7, 543–570 (2021).

5. Grossman, S. et al. Convergent evolution of face spaces across human face-selective neuronal groups and deep convolutional networks. Nat. Commun. 10, 1–13 (2019).

6. Dobs, K., Martinez, J., Kell, A. J. E. & Kanwisher, N. Brain-like functional specialization emerges spontaneously in deep neural networks. Sci. Adv. 8, eabl8913 (2022).

7. Abudarham, N., Grosbard, I. & Yovel, G. Face recognition depends on specialized mechanisms tuned to view-invariant facial features: Insights from deep neural networks optimized for face or object recognition. Cogn. Sci. 45, e13031 (2021).

8. Jacob, G., Pramod, R. T., Katti, H. & Arun, S. P. Qualitative similarities and differences in visual object representations between brains and deep networks. Nat. Commun. 12, 1–14 (2021).

9. Yovel, G., Grosbard, I. & Abudarham, N. Testing the Expertise Hypothesis with Deep Convolutional Neural Networks Optimized for Subordinate-level Categorization. VSS Conf. (2022).

10. Tian, F., Xie, H., Song, Y., Hu, S. & Liu, J. The Face Inversion Effect in Deep Convolutional Neural Networks. Front. Comput. Neurosci. 16, (2022).

11. Cavazos, J. G., Jeckeln, G., Hu, Y. & O’Toole, A. J. Strategies of Face Recognition by Humans and Machines. in Deep Learning-Based Face Analytics 361–379 (Springer, 2021).

12. Rademacher, L. et al. Dissociation of neural networks for anticipation and consumption of monetary and social rewards. Neuroimage 49, 3276–3285 (2010).

13. Lohr, S. Facial recognition is accurate, if you’re a white guy. in Ethics of Data and Analytics 143–147 (Auerbach Publications, 2018).

14. Hugenberg, K., Young, S. G., Bernstein, M. J. & Sacco, D. F. The Categorization-Individuation Model: An Integrative Account of the Other-Race Recognition Deficit. Psychol. Rev. 117, 1168–1187 (2010).

15. Meissner, C. A. & Brigham, J. C. Thirty years of investigating the own-race bias in memory for faces: A meta-analytic review. Psychol. Public Policy, Law 7, 3 (2001).

16. Yovel, G. & Abudarham, N. From concepts to percepts in human and machine face recognition: A reply to Blauch, Behrmann \& Plaut. Cognition 208, 104424 (2021).

17. Young, A. W. & Burton, A. M. Are we face experts? Trends Cogn. Sci. 22, 100–110 (2018).

18. Blauch, N. M., Behrmann, M. & Plaut, D. C. Deep learning of shared perceptual representations for familiar and unfamiliar faces: Reply to commentaries. Cognition 208, 104484 (2021).

19. Noyes, E. et al. Seeing through disguise: Getting to know you with a deep convolutional neural network. Cognition 211, 104611 (2021).

20. Gobbini, M. I. & Haxby, J. V. Neural systems for recognition of familiar faces. Neuropsychologia 45, 32–41 (2007).

21. Bruce, V. & Young, A. Understanding face recognition. Br. J. Psychol. 77, 305–327 (1986).

22. Schwartz, L. & Yovel, G. The roles of perceptual and conceptual information in face recognition. J. Exp. Psychol. Gen. 145, 1493–1511 (2016).

23. Radford, A. et al. Learning Transferable Visual Models From Natural Language Supervision. (2021) doi:10.48550/arxiv.2103.00020.

24. Muennighoff, N. SGPT: GPT Sentence Embeddings for Semantic Search. arXiv Prepr. (2022).

25. Simonyan, K. & Zisserman, A. Very deep convolutional networks for large-scale image recognition. arXiv Prepr. arXiv1409.1556 (2014).

26. der Maaten, L. & Hinton, G. Visualizing data using t-SNE. J. Mach. Learn. Res. 9, (2008).

27. Tsantani, M. et al. Ffa and ofa encode distinct types of face identity information. J. Neurosci. 41, 1952–1969 (2021).

28. Yildirim, I., Belledonne, M., Freiwald, W. & Tenenbaum, J. Efficient inverse graphics in biological face processing. Sci. Adv. 6, eaax5979 (2020).

29. Jozwik, K. M. et al. Face dissimilarity judgments are predicted by representational distance in morphable and image-computable models. Proc. Natl. Acad. Sci. 119, e2115047119 (2022).

30. Song, Y., Qu, Y., Xu, S. & Liu, J. Implementation-independent representation for deep convolutional neural networks and humans in processing faces. Front. Comput. Neurosci. 14, 601314 (2021).

31. Young, A. W. & Bruce, V. Understanding person perception. Br. J. Psychol. 102, 959–974 (2011).

32. Karras, T., Laine, S. & Aila, T. A Style-Based Generator Architecture for Generative Adversarial Networks. IEEE Trans. Pattern Anal. Mach. Intell. 43, 4217–4228 (2018).

33. Karras, T. et al. Analyzing and Improving the Image Quality of StyleGAN. Proc. IEEE Comput. Soc. Conf. Comput. Vis. Pattern Recognit. 8107–8116 (2019) doi:10.1109/CVPR42600.2020.00813.

34. Clarke, A. & Tyler, L. K. Object-specific semantic coding in human perirhinal cortex. J. Neurosci. 34, 4766–4775 (2014).

35. Landi, S. M. & Freiwald, W. A. Two areas for familiar face recognition in the primate brain. Science (80-.). 357, 591–595 (2017).

36. Landi, S. M., Viswanathan, P., Serene, S. & Freiwald, W. A. A fast link between face perception and memory in the temporal pole. Science (80-.). 373, 581–585 (2021).

37. Parde, C. J. et al. Twin identification over viewpoint change: A deep convolutional neural network surpasses humans. arXiv Prepr. arXiv2207.05316 (2022).

38. Radford, A. et al. Learning Transferable Visual Models From Natural Language Supervision. (2021).

39. Cao, Q., Shen, L., Xie, W., Parkhi, O. M. & Zisserman, A. VGGFace2: A dataset for recognising faces across pose and age. Proc. - 13th IEEE Int. Conf. Autom. Face Gesture Recognition, FG 2018 67–74 (2017) doi:10.48550/arxiv.1710.08092.

40. Zhang, K., Zhang, Z., Li, Z. & Qiao, Y. Joint Face Detection and Alignment using Multitask Cascaded Convolutional Networks. IEEE Signal Process. Lett. 23, 1499–1503 (2016).

41. Parkhi, O. M., Vedaldi, A. & Zisserman, A. Deep face recognition. BMVC 2015 - Proc. Br. Mach. Vis. Conf. 2015 (2015).

42. Huang, G. B., Ramesh, M., Berg, T. & Learned-Miller, E. Labeled Faces in the Wild: A Database for Studying Face Recognition in Unconstrained Environments. ICCV (2007).

43. Ma, N., Baetens, K., Vandekerckhove, M., Van der Cruyssen, L. & Van Overwalle, F. Dissociation of a trait and a valence representation in the mPFC. Soc. Cogn. Affect. Neurosci. 9, 1506–1514 (2013).

