## supplementary data for "Deep learning algorithms reveal a new visual-semantic representation of familiar faces in human perception and memory"

**Supplementary information**

Supplementary notes include additional information about results and procedures

1. Semantic similarity rating

To estimate the semantic similarity based on humans' ratings, participants were asked to rate the semantic similarity of all identity pairs based on their images or their names. In both cases, participants were asked to ignore the visual appearance of the identities and only rate them based on their biographical information. The correlation between human semantic ratings based on pictures and names were very high (r= 0.94) indicating that either of them can be used to measure human semantic similarity. In Figure 4 in the main text, we display the correlations of the DNNs based on the semantic name similarity task. Supplementary Figures 5 and 6 also show the correlations of the three DNNs with human semantic ratings, showing a very high correlation between SGPT and human semantic ratings.

2. CLIP and VGG generated images:

Training a model to generate StyleGAN's embeddings:

To train this model, we used a subset of images from the CelebA-HQ dataset [1], all of which were identities familiar to CLIP (tested with zero-shot classification of name-face images described in the methods section), and none of them depict identities used in the main experiment. We mapped the embeddings of these images according to VGG and CLIP to StyleGAN's embeddings. To obtain this mapping we trained this model in the following steps:

- Each image embedding was paired with the StyleGAN embedding describing the same image, which was calculated in a process termed StyleGAN inversion [2] by using the e4e algorithm [3].
- These image pairs were used to train the model to transform a DNN (VGG or CLIP) embedding to a StyleGAN embedding. We trained 18 independent normalizing

flows using the RealNVP architecture [4], mapping between the 512-dimensional latent variable of each input layer of StyleGAN [5] , conditioned on the image's representation according to each DNN, to the 512-dimensional multivariate normal distribution.

- All RealNVP models were optimized using the Adam optimization algorithm [6] with a learning rate of 1e-5 and default PyTorch [7] parameters for 400,000 iterations with batch size of 12 images. Learning rate reduced by a factor of 10 after 200,000 iterations. Image reconstruction from CLIP embeddings was done with representations from the ViT-B/32 architecture [8].

We ran a similar procedure with a model that was trained on different datasets. We used CelebAHQ full dataset, VGGFace2 dataset (familiar to CLIP) or CelebA full dataset. We trained the model between StyleGAN and CLIP using each of these datasets and the procedure described above. To test if the different trained models generated similar images we generated an RDM using VGG-16 for each set and calculated the correlations between these RDMs. The correlations were very high (r = 0.89-95), indicating that the models generated similar representations. Supplementary Figure 7 shows the generated images based on CLIP and VGG models.

Recognition of VGG and CLIP-generated faces:

To test if the generated images can be recognized by humans, we performed a two-stage face recognition task: Participants either were assigned to recognize VGG-generated faces or CLIP-generate faces. The task included two stages: a free recall recognition and a face-name matching from a list of names. All the participants conducted the free recall task prior to the face-name matching task. In the free-recall task, the participants were presented with a set of face images and for each image, they had to indicate to which familiar person the face is mostly similar. The participants could also write a description of this person if they could not remember their name or to indicate that they do not recognize the face. Following the free recall, the participants performed face name matching. In this task, they were presented with the same set of face images, and each image was presented with a list of names of highly famous identities. They were asked to indicate for each face, to which of the identities in the list, the face is most similar. All female faces were presented with the same 18 names (9 names of faces that were included in our task and 9 names of novel identities) and all male faces were presented with the same 22 names (11 names of faces that were included in our task and 11 names of novel identities). The full list of names in shown in Supplementary Figure 9. After they completed both recognition tasks, the participants were asked to indicate for the original 20 identities images whether they were familiar with them before the experiment. The experiment lasted about 15 minutes.

We excluded from the analysis trials that included generated images that are based on identities that the participant was not familiar with. In addition, in a few cases participants indicated that they are not familiar with an identity based on its original image, but did match the face to the correct name or recognize them correctly based on the DNN-generated image. In such cases we marked them as a correct response and included these trials in the analysis.

We first assessed whether the VGG- and CLIP-generated faces can be recognized as the original identities by humans. Participants were presented with each VGG- or CLIP-generated face and were asked to write their name or if they do not recall the name to write any unique semantic information that they know about them. We computed the proportion of participants who correctly recognized each face image. The VGG-generated faces were recognized by an average of 51% (median: 59%; range: 0-90%) of the participants across the 20 face images, and CLIP-generated faces were recognized by an average of 31% (median: 28%; range: 0-60%) of the participants across the 20 face images. To assess whether performance is improved in a face-name matching task, in a second phase of the experiment participants were presented with a list of different names of celebrities from the same gender (18 females and 22 males) and were asked to match the face image to the name of the person that is most similar to them. The correct name was selected by the majority of the participants for 18/20 images in VGG-generated set and for 19/20 in CLIP-generated set. The VGG-generated faces were correctly matched to their names by an average of 76% (median: 84%; range: 5-100%) of the participants across the 20 face images, and CLIP-generated faces were correctly matched to their names by an average of 57% (median: 57%; range: 0-95%) of the participants across the 20 face images. The results of this analysis for each identity are reported in supplementary Figure 10. These results show that the VGG- and CLIP-generated faces were overall well recognized.

Supplementary figures:

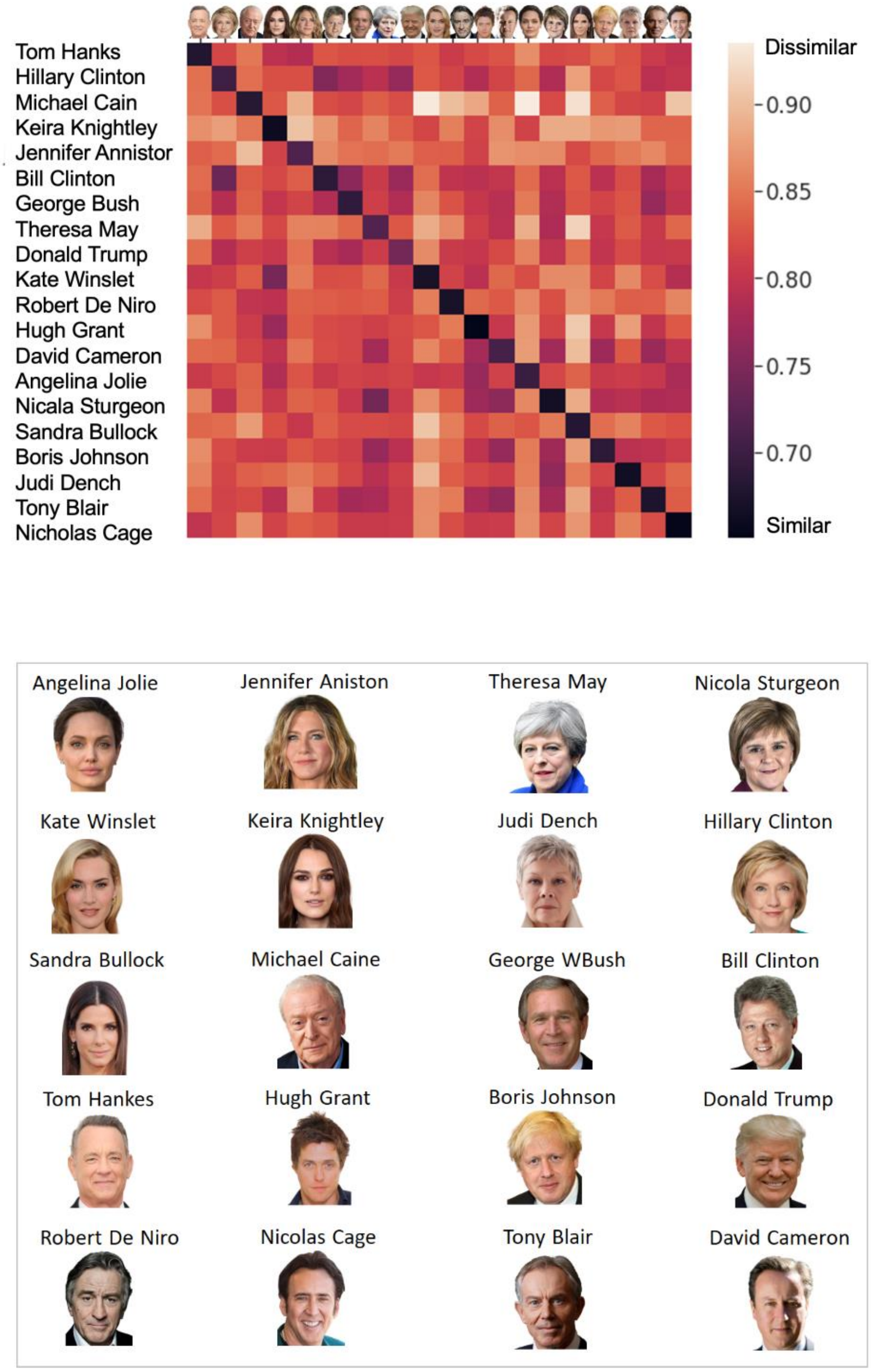


Supplementary Figure 1: Top: RDM of the CLIP embeddings of face images and names. Each name was most similar to the corresponding face image than any other identity (diagonal). Bottom: The images of the faces of the 20 celebrities were selected from Google images.

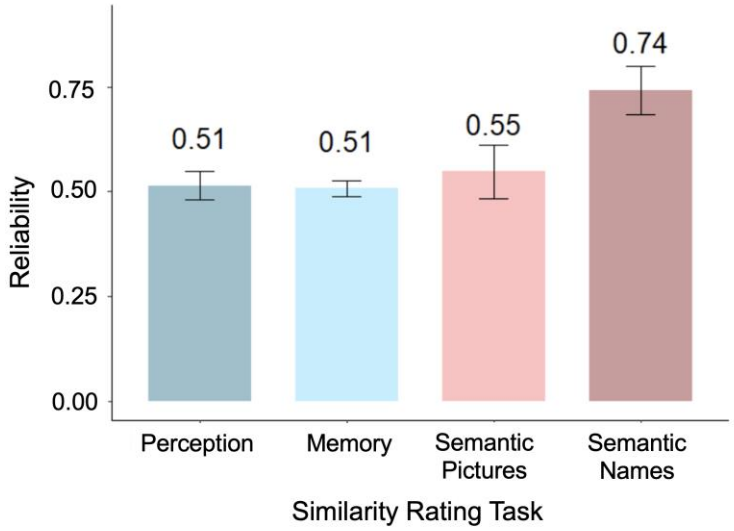


Supplementary Figure 2: The reliability of human similarity ratings in the visual and semantic tasks was assessed by calculating the Pearson correlation of the RDM of each participant with the average RDM across all other participants (leave-one out). The figure shows the averaged correlation of all leave one out correlations. The error bar indicates the standard deviation.

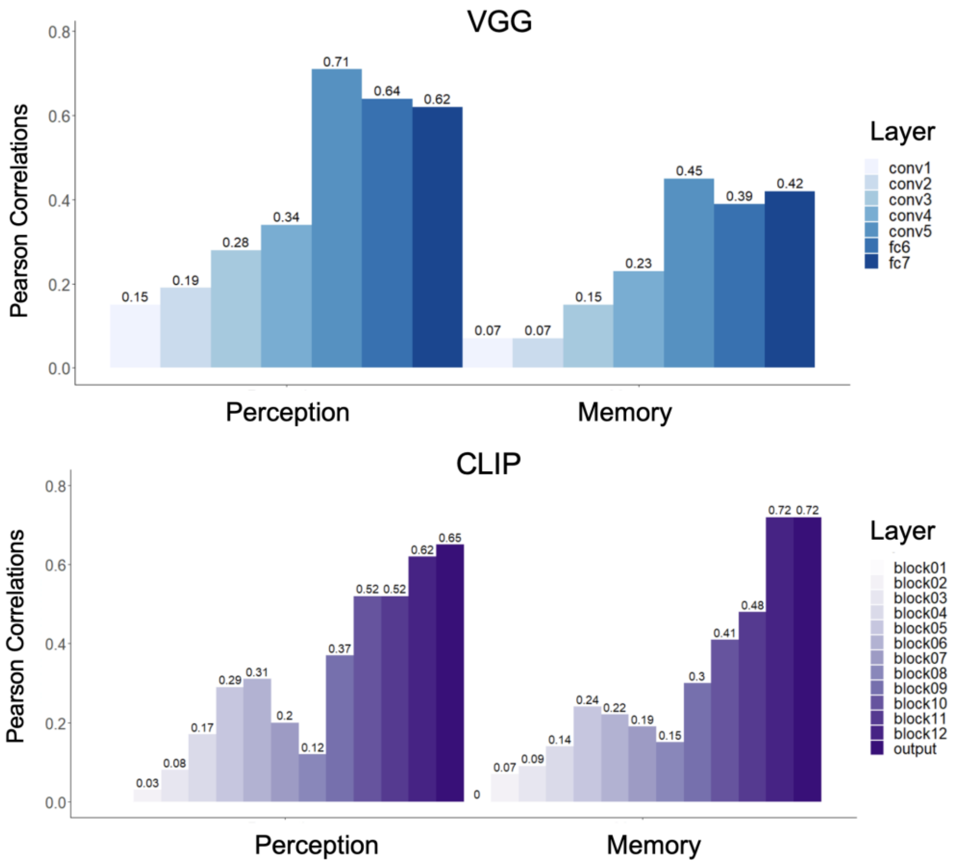


Supplementary Figure 3: The correlations between similarity scores for all pairs of faces based on the embeddings of each layer of VGG-16 (Top) and each layer of CLIP (Bottom) with human visual similarity ratings in perception (left) and memory (right). Correlations between DNN and human representations were lower in lower layers and increased for higher layers.

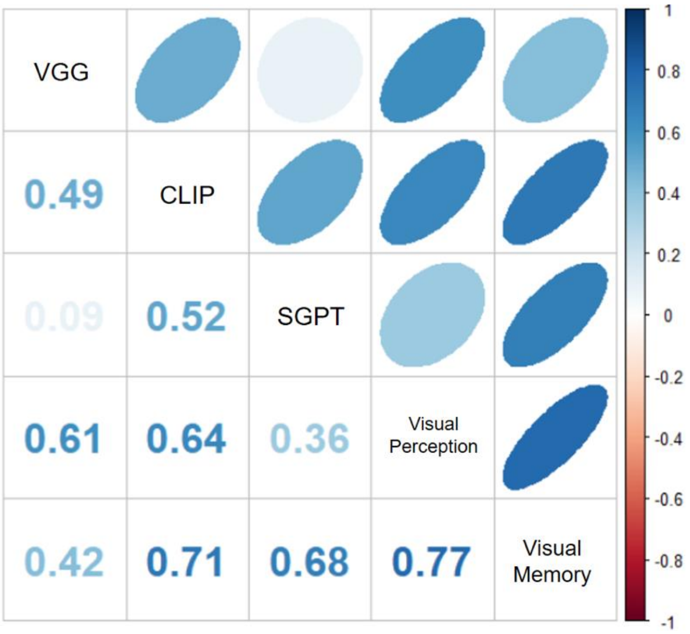


Supplementary Figure 4: The correlations between the RDMs of the DNNs (VGG, CLIP. SGPT) and human visual similarity ratings in perception and memory of the same face identities.

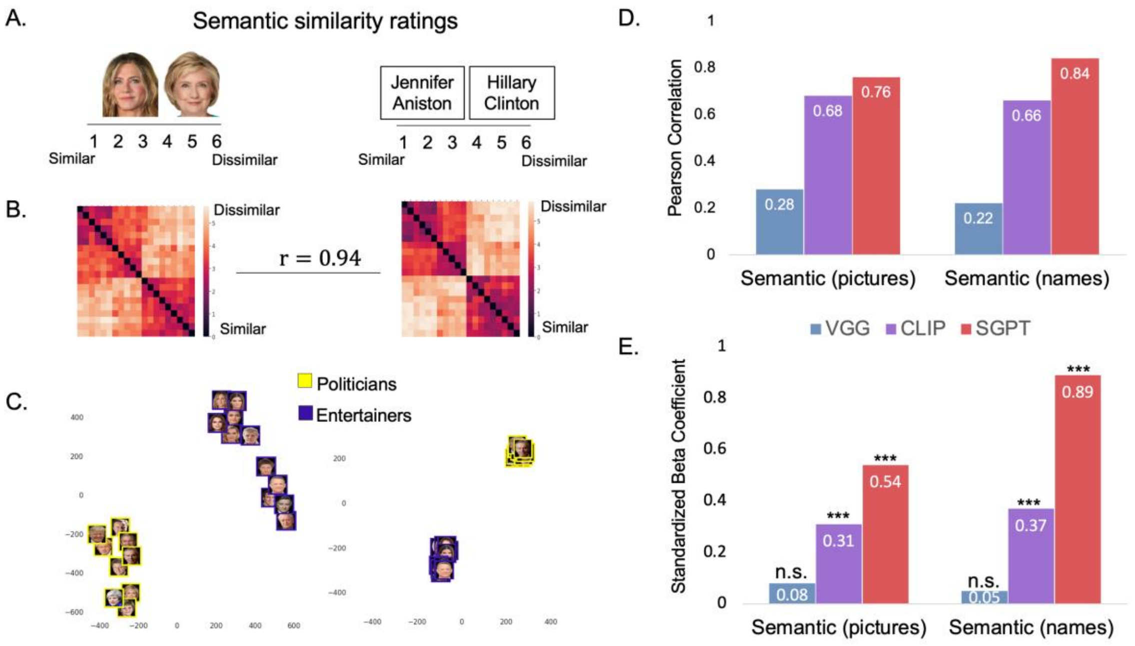


Supplementary Figure 5: A. Participants rated the semantic similarity between familiar identities when they were presented with pictures or their names. B. The RDMs based on semantic ratings ordered by occupation. The RDMs of the two tasks were highly correlated. C. A t-SNE visualization of the RDMs, showing semantic influence according to occupation. D. Correlations between the RDMs based on embeddings of the same identities in visual (VGG), visual-semantic (CLIP) and semantic (SGPT) DNNs with human semantic representations. E. Results of the multiple linear regression model with the three DNNs as predictors of human semantic ratings show that for both semantic ratings tasks, SGPT and CLIP significantly predicted human semantic representations, whereas VGG did not.

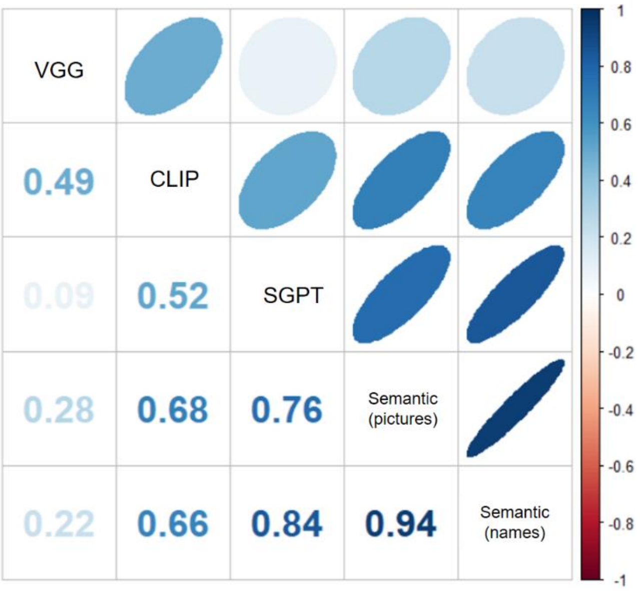


Supplementary Figure 6: Correlation matrix between human semantic RDMs (based on pictures and names) and the visual (VGG), visual semantic (CLIP) and the semantic (SGPT) DNNs.

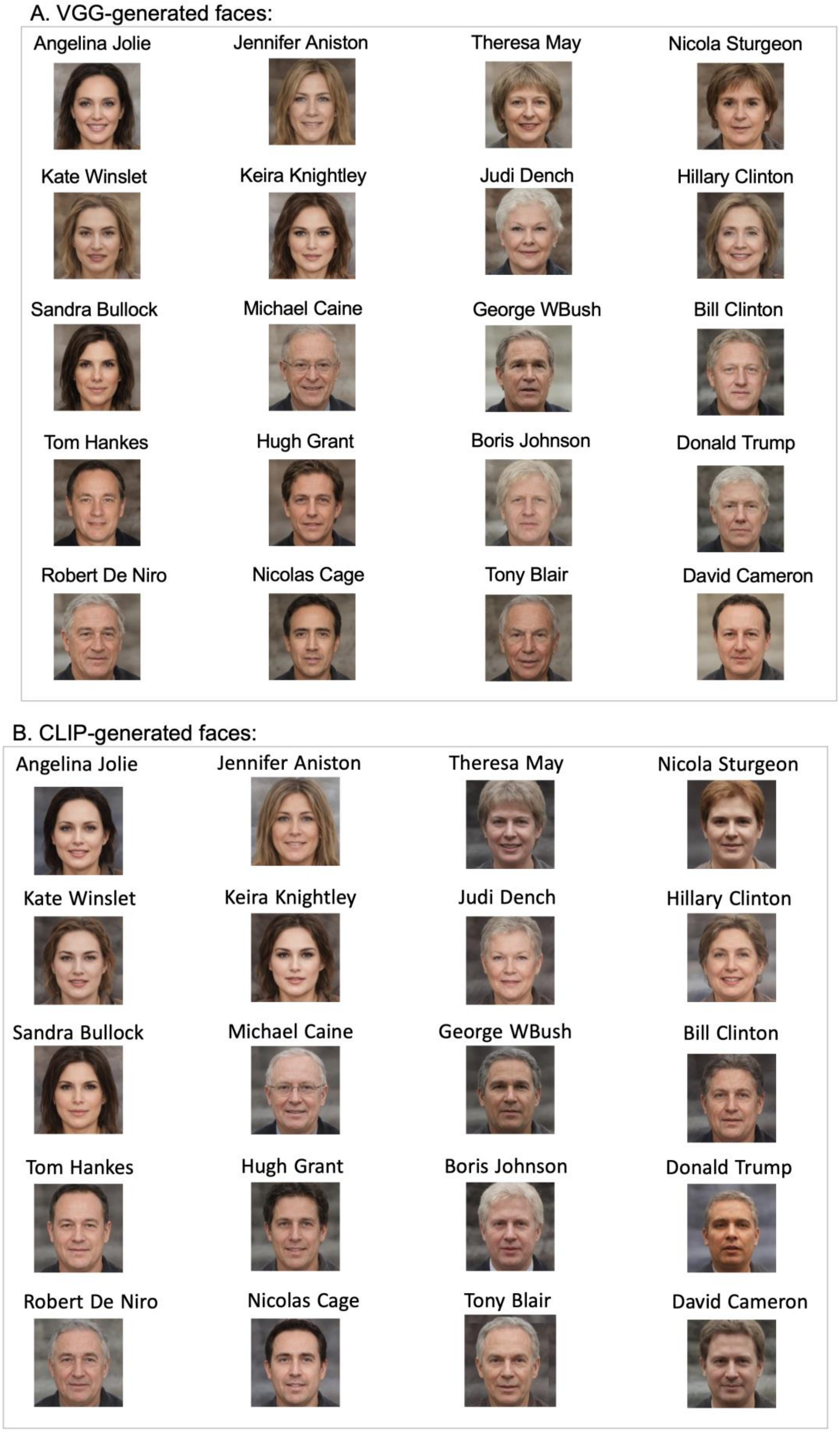


Supplementary Figure 7: The images generated by StyleGAN based on the averaged embeddings of 20 different images of each identity based on A. VGG-16. B. CLIP

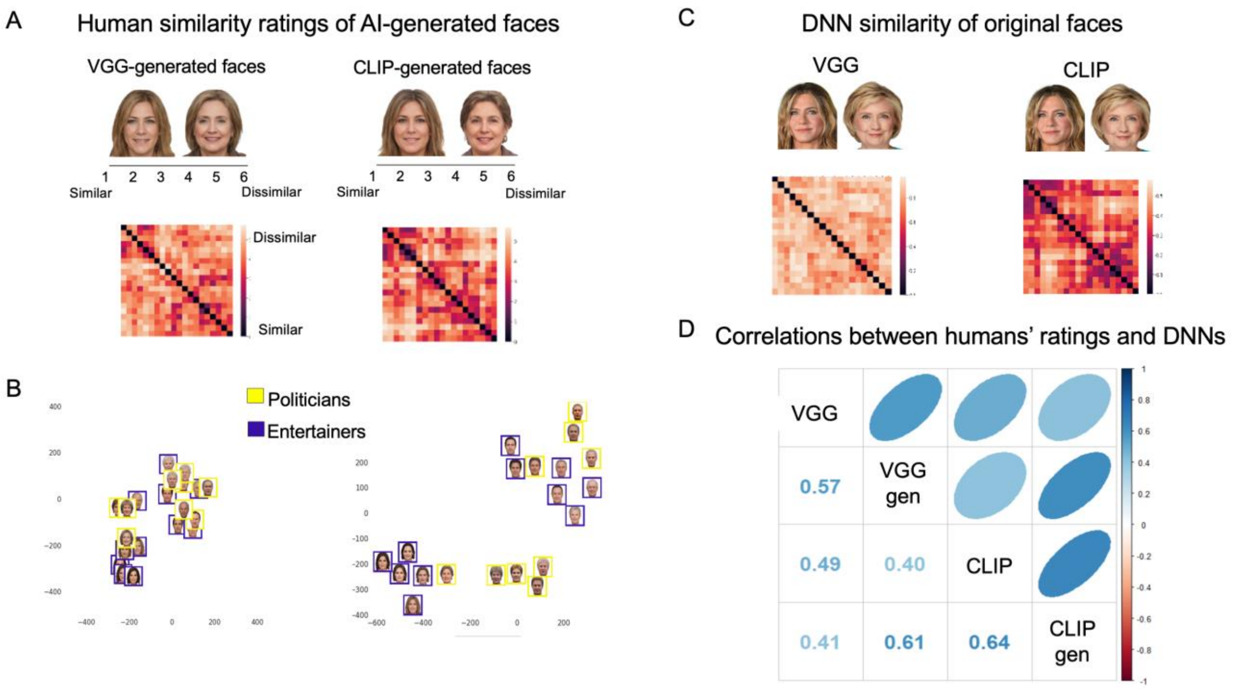


Supplementary Figure 8: A. Participants rated the visual similarity between VGG- and CLIP-generated faces. RDM shows human similarity scores B. A t-SNE visualization of human RDMs C. CLIP and VGG RDMs based on the distance between embeddings of the original images.D. Correlations between human similarity ratings of VGG and CLIP-generated faces and DNNs similarity measures of the original faces: VGG: VGG embeddings; VGG-gen: human similarity ratings of the VGG-generated faces. CLIP: CLIP embeddings; CLIP-gen: human similarity ratings of the CLIP-generated faces.

| Female names: | Male names: |
|---|---|
| Angela Merkel | Arnold Schwarzenegger |
| Angelina Jolie | Bill Clinton |
| Hillary Clinton | Boris Johnson |
| Jennifer Aniston | Clint Eastwood |
| Judi Dench | David Cameron |
| Julia Roberts | Donald Trump |
| Kate Middleton | George Clooney |
| Kate Winslet | George W Bush |
| Keira Knightley | Hugh Grant |
| Meryl Streep | Joe Biden |
| Nicola Sturgeon | Leonardo Dicaprio |
| Nicole Kidman | Michael Caine |
| Penelope Cruise | Nicola Sarkozy |
| Queen Elizabeth | Nicolas Cage |
| Reese Witherspoon | Prince William |
| Sandra Bullock | Richard Gere |
| Sarah Jessica Parker | Robert De Niro |
| Theresa May | Sylvester Stallone |
| | Tom Cruise |
| | Tom Hanks |
| | Tony Blair |

Supplementary Figure 9: Female and male names used in the face-name matching task with the VGG- and CLIP-generated faces.

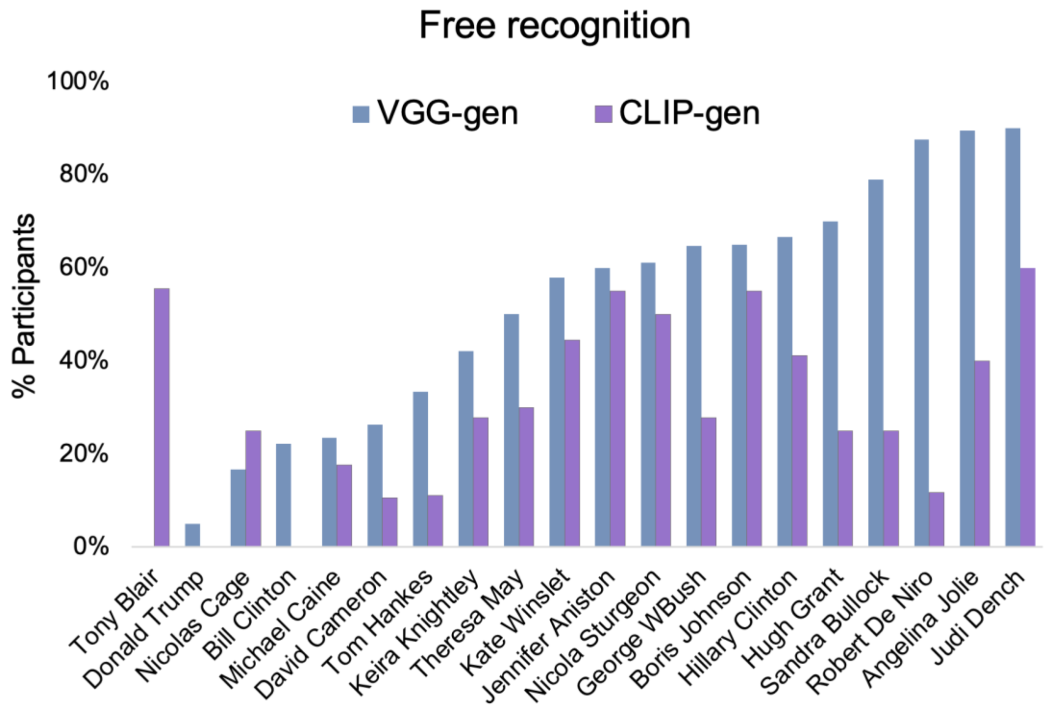


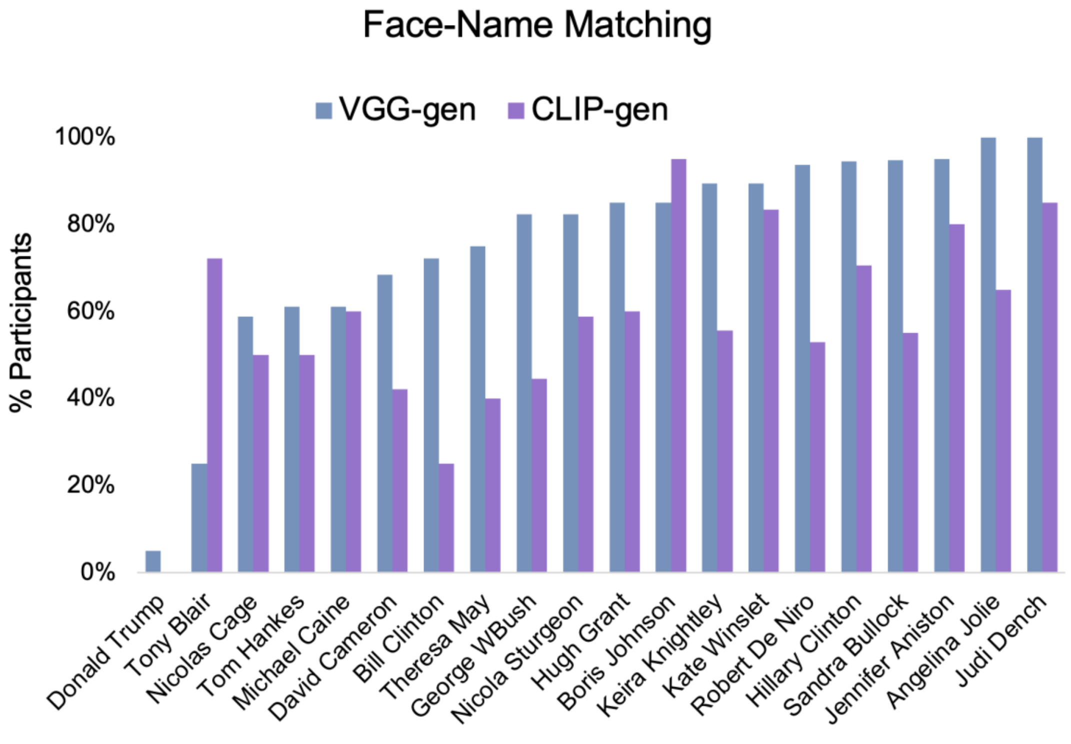


Supplementary Figure 10: The percentage of participants who recognized each of the identities based on the VGG and CLIP-generated faces. Top: in the face recognition task. Bottom: in a face-name matching task.

Supplementary Table 1: The first paragraph in Wikipedia of each identity was used to extract embeddings of the semantic representations of the 20 identities from SGPT:

| Identitiy | Text |
|---|---|
| Angelina Jolie | Angelina Jolie[3] DCMG (born Angelina Jolie Voight,[4] June 4, 1975; later Angelina Jolie Pitt[5]) is an American actress, filmmaker, and humanitarian. The recipient of numerous accolades, including an Academy Award and three Golden Globe Awards, she has been named Hollywood's highest-paid actress multiple times. |
| Jennifer Aniston | Jennifer Joanna Aniston (born February 11, 1969) is an American actress and producer. The daughter of actors John Aniston and Nancy Dow, she began working as an actress at an early age with an uncredited role in the 1988 film Mac and Me; her first major film role came in the 1993 horror comedy Leprechaun. Since her career progressed in the 1990s, she has become one of the worlds highest-paid actresses. |
| Judi Dench | Dame Judith Olivia Dench CH DBE FRSA (born 9 December 1934) is an English actress. Regarded as one of Britain's best actresses,[1][2][3] she is noted for her versatile work in various films and television programmes encompassing several genres, as well as for her numerous roles on the stage.[4] Dench has garnered various accolades throughout her career spanning over six decades, including an Academy Award, a Tony Award, two Golden Globe Awards, four British Academy Television Awards, six British Academy Film Awards and seven Olivier Awards. |
| Kate Winslet | Kate Elizabeth Winslet CBE (born 5 October 1975) is an English actress.[3] Known for her work in independent films, particularly period dramas, and for her portrayals of headstrong and complicated women, she has received numerous accolades, including an Academy Award, a Grammy Award, two Primetime Emmy Awards, three British Academy Film Awards, and five Golden Globe Awards. Time magazine named Winslet one of the 100 most influential people in the world in 2009 and 2021, and in 2012, she was appointed Commander of the Order of the British Empire (CBE). |
| Keira Knightley | Keira Christina Righton[1] OBE Knightley, born 26 March 1985) is an English actress.[2] She has starred in both independent films and big-budget blockbusters, and is particularly noted for her roles in period dramas. Her accolades include two Empire Awards and nominations for two Academy Awards, three British Academy Film Awards, three Golden Globe Awards, one Screen Actors Guild Award and one Laurence Olivier Award. Knightley was appointed an OBE in the 2018 Birthday Honours for services to drama and charity.[3] |
| Sandra Bullock | Sandra Annette Bullock ( born July 26, 1964) is an American actress and producer. The recipient of various accolades, including an Academy Award and a Golden Globe Award, she was the world's highest-paid actress in both 2010 and 2014.[1][2][3] In 2010, she was named one of Time's 100 most influential people in the world. |

| | |
|---|---|
| Hugh Grant | Hugh John Mungo Grant[2] (born 9 September 1960) is an English actor. His awards include a Golden Globe Award, a BAFTA Award, Volpi Cup and an Honorary C?sar. As of 2018, his films have grossed a total of nearly US$3 billion worldwide from 29 theatrical releases.[3] |
| Michael Caine | Sir Michael Caine CBE (born Maurice Joseph Micklewhite; 14 March 1933) is an English actor. Known for his distinctive South London accent, he has appeared in more than 160 films in a career spanning seven decades, and is considered a British film icon.[2][3] He has received various awards including two Academy Awards, a BAFTA Award, three Golden Globe Awards, and a Screen Actors Guild Award. As of February 2017, the films in which Caine has appeared have grossed over $7.8 billion worldwide.[4] Caine is one of only five male actors to be nominated for an Academy Award for acting in five different decades.[nb 1] He has appeared in seven films that featured in the British Film Institute's 100 greatest British films of the 20th century. In 2000, he received a BAFTA Fellowship and was knighted by Queen Elizabeth II for his contribution to cinema. |
| Nicolas Cage | Nicolas Kim Coppola (born January 7, 1964),[2][3] known professionally as Nicolas Cage, is an American actor and filmmaker. Born into the Coppola family, Cage is the recipient of various accolades, including an Academy Award, a Screen Actors Guild Award, and a Golden Globe Award. |
| Robert De Niro | Robert Anthony De Niro Jr. (/d? ?n??ro?/ d? NEER-oh, Italian: [de ?ni?ro]; born August 17, 1943) is an American actor, producer, and director. He is particularly known for his nine collaborations with filmmaker Martin Scorsese, and is the recipient of various accolades, including two Academy Awards, a Golden Globe Award, the Cecil B. DeMille Award, and a Screen Actors Guild Life Achievement Award. In 2009, De Niro received the Kennedy Center Honor, and received a Presidential Medal of Freedom from U.S. President Barack Obama in 2016. |
| Tom Hankes | Thomas Jeffrey Hanks (born July 9, 1956) is an American actor and filmmaker. Known for both his comedic and dramatic roles, he is one of the most popular and recognizable film stars worldwide, and is regarded as an American cultural icon.[2] Hanks's films have grossed more than $4.9 billion in North America and more than $9.96 billion worldwide,[3] making him the fourth-highest-grossing actor in North America.[4] |
| Hillary Clinton | Hillary Diane Rodham Clinton (born October 26, 1947) is an American politician, diplomat, lawyer, writer, and public speaker who served as the 67th United States secretary of state from 2009 to 2013, as a United States senator representing New York from 2001 to 2009, and as first lady of the United States from 1993 to 2001 as the wife of President Bill Clinton. A member of the Democratic Party, she was the party's nominee for president in the 2016 presidential election, which she lost to Donald Trump. |
| Nicola Sturgeon | Nicola Ferguson Sturgeon (born 19 July 1970) is a Scottish lawyer and politician serving as First Minister of Scotland and Leader of the Scottish National Party (SNP) since 2014. She is the first woman to hold either position. She has been a member of the Scottish Parliament (MSP) since 1999, first as an additional member for the Glasgow electoral region, and as the member for Glasgow Southside (formerly Glasgow Govan) from 2007. |
| Theresa May | Theresa Mary, Lady May[1] (Brasier; born 1 October 1956) is a British politician who served as Prime Minister of the United Kingdom and Leader of the Conservative Party from 2016 to 2019. She served as Home Secretary from 2010 to 2016 in the Cameron government and has been the |

| | |
|---|---|
| | Member of Parliament (MP) for Maidenhead in Berkshire since 1997. Ideologically, May identifies herself as a one-nation conservative.[3] |
| Bill Clinton | William Jefferson Clinton ( Blythe III; born August 19, 1946) is an American politician who served as the 42nd president of the United States from 1993 to 2001. He previously served as governor of Arkansas from 1979 to 1981 and again from 1983 to 1992, and as attorney general of Arkansas from 1977 to 1979. A member of the Democratic Party, Clinton became known as a New Democrat, as many of his policies reflected a centrist "Third Way" political philosophy. He is the husband of Hillary Clinton, who was a senator from New York from 2001 to 2009, secretary of state from 2009 to 2013 and the Democratic nominee for president in the 2016 presidential election. |
| Boris Johnson | Alexander Boris de Pfeffel Johnson (born 19 June 1964) is a British politician serving as Prime Minister of the United Kingdom and Leader of the Conservative Party since 2019. He was Secretary of State for Foreign and Commonwealth Affairs from 2016 to 2018 and Mayor of London from 2008 to 2016. Johnson has been Member of Parliament (MP) for Uxbridge and South Ruislip since 2015 and was previously MP for Henley from 2001 to 2008. |
| David Cameron | David William Donald Cameron (born 9 October 1966) is a British politician, businessman, lobbyist, and author who served as Prime Minister of the United Kingdom from 2010 to 2016. He was Member of Parliament (MP) for Witney from 2001 to 2016 and leader of the Conservative Party from 2005 to 2016. He identifies as a one-nation conservative, and has been associated with both economically liberal and socially liberal policies. |
| Donald Trump | Donald John Trump (born June 14, 1946) is an American politician, media personality, and businessman who served as the 45th president of the United States from 2017 to 2021. |
| George Bush | George Walker Bush (born July 6, 1946) is an American politician who served as the 43rd president of the United States from 2001 to 2009. A member of the Bush family and son of former president George H. W. Bush, he previously served as the 46th governor of Texas from 1995 to 2000 as part of the Republican Party. |
| Tony Blair | Sir Anthony Charles Lynton Blair KG (born 6 May 1953) is a British politician who served as Prime Minister of the United Kingdom from 1997 to 2007 and Leader of the Labour Party from 1994 to 2007. On his resignation he was appointed Special Envoy of the Quartet on the Middle East, a diplomatic post which he held until 2015. He has been the executive chairman of the Tony Blair Institute for Global Change since 2016. As prime minister, many of his policies reflected a centrist "Third Way" political philosophy.[b] He is the only living former Labour leader to have led the party to a general election victory; and one of only two in history to form three majority governments, the other being Harold Wilson. |

References:

1. Karras, T., Aila, T., Laine, S. & Lehtinen, J. Progressive Growing of GANs for Improved Quality, Stability, and Variation. *6th Int. Conf. Learn. Represent. ICLR 2018 - Conf. Track Proc.* (2017) doi:10.48550/arxiv.1710.10196.

2. Xia, W. *et al.* Gan inversion: A survey. *IEEE Trans. Pattern Anal. Mach. Intell.* (2022).

3. Tov, O., Alaluf, Y., Nitzan, Y., Patashnik, O. & Cohen-Or, D. Designing an encoder for StyleGAN image manipulation. *ACM Trans. Graph.* (2021) doi:10.1145/3450626.3459838.

4. Dinh, L., Sohl-Dickstein, J. & Bengio, S. Density estimation using Real NVP. *5th Int. Conf. Learn. Represent. ICLR 2017 - Conf. Track Proc.* (2016) doi:10.48550/arxiv.1605.08803.

5. Abdal, R., Qin, Y. & Wonka, P. Image2StyleGAN: How to Embed Images Into the StyleGAN Latent Space? *Proc. IEEE Int. Conf. Comput. Vis.* **2019-October**, 4431–4440 (2019).

6. Kingma, D. P. & Ba, J. L. Adam: A Method for Stochastic Optimization. *3rd Int. Conf. Learn. Represent. ICLR 2015 - Conf. Track Proc.* (2014) doi:10.48550/arxiv.1412.6980.

7. Paszke, A. *et al.* PyTorch: An Imperative Style, High-Performance Deep Learning Library. *Adv. Neural Inf. Process. Syst.* **32**, (2019).

8. Radford, A. *et al.* Learning Transferable Visual Models From Natural Language Supervision. (2021).

.
